# Comparative analysis of seed size, germination, and vegetative allocation in annual and herbaceous perennial crops and their wild relatives in *Lupinus* and *Phaseolus* (Fabaceae)

**DOI:** 10.1101/824375

**Authors:** Sterling A. Herron, Matthew J. Rubin, Claudia Ciotir, Timothy E. Crews, David L. Van Tassel, Allison J. Miller

## Abstract

Herbaceous perennial species are receiving increased attention for their potential to provide both edible products and ecosystem services in agricultural systems. Many legumes (Fabaceae Lindl.) are of special interest due to nitrogen fixation carried out by bacteria in their roots and their production of protein-rich, edible seeds. However, herbaceous perennial legumes have yet to enter widespread use as pulse crops, and the response of wild, herbaceous, perennial species to artificial selection for increased seed yield remains under investigation. Here we compare cultivated and wild accessions of congeneric annual and herbaceous perennial legume species to investigate associations of lifespan and cultivation with seed size, germination, and first year vegetative growth patterns, and to characterize covariation among traits. We use “cultivated” to describe accessions with a history of human planting and use, which encompasses a continuum of domestication. Analyses focused on three annual and eight perennial *Lupinus* species, and three annual and four perennial *Phaseolus* species. We found a significant association of both lifespan and cultivation status with seed size (weight, area, length), germination proportion, node number, stem diameter, shoot dry mass, and root dry mass. Wild seed size was greater in annuals for *Lupinus* and greater for perennials in *Phaseolus*. Germination was lower in wild perennials than wild annuals in both genera, and vegetative allocation was roughly equivalent across lifespans in wild *Phaseolus*. Relative to wild forms, both cultivated annual and cultivated perennial accessions exhibited greater seed size, lower germination proportion, and larger overall plant size. Seed size traits were positively correlated with vegetative growth traits, and all biomass traits examined here were positively correlated. This study highlights some basic similarities and differences between annual and herbaceous perennial legumes, and provides insights into how perennial legumes might respond to artificial selection compared to annual species.

## 1 INTRODUCTION

At least one grass (Poaceae) species and one legume (Fabaceae) species were domesticated in most centers of domestication (Vavilov, 1992). However nearly all grass and herbaceous legume crops grown for human consumption are annual plants that complete their life cycle in a single year, or are perennial species cultivated as annuals (Van Tassel et al., 2010). Although thousands of herbaceous perennial species exist within major crop families (e.g., Ciotir et al., 2016; Ciotir et al., 2019), herbaceous annual plant species were likely initially selected during the early stages of domestication due to pre-existing agriculturally favorable traits, e.g., high reproductive yield in a single season and accelerated germination and flowering (Van Tassel et al., 2010). Over time, artificial selection has led to exceptional gains in seed production in annual crops, particularly in the last century (Mann, 1997); however, cultivation intensity and other agronomic practices have resulted in widespread soil loss (FAO, 2019). Current research focuses in part on the development of multi-functional crops to support the ecological intensification of agriculture, which targets high yields as well as ecosystem services, such as soil and water retention (Ryan et al., 2018).

Herbaceous perennial species have been proposed as key components of ecological intensification (Jordan & Warner, 2010; Ryan et al., 2018). Longer-lived species have deep, persistent root systems that mitigate erosion and enhance nutrient uptake, they produce perennating shoots which reduce yearly planting costs, and have a longer photosynthetically active growth period, allowing for high biomass production each year (Cox et al., 2006; Crews et al., 2018). Due to their ability to tolerate stressful, resource-poor conditions, herbaceous perennial crops also have applications in arid lands unsuitable for annuals (Cox et al., 2006). Further, multiple perennial species grown together in polyculture have the potential to increase pest resistance, inhibit weeds, and foster a diverse and healthy soil microbiome (Crews et al., 2018). However, only a few perennial seed crops (principally cereals, oilseeds, and pulses) have entered the domestication process, and we know relatively little about how artificial selection for increased seed production will impact the rest of the perennial plant.

Life history theory has long been used as a framework for understanding differences in survival, reproduction, and resource allocation in annual and perennial plants (Cole, 1954; Charnov & Schaffer, 1973). Annual and perennial species are largely thought to be at opposite ends of the plant economics spectrum, one of several widely used concepts within life history theory (Reich, 2014). In this model, plants either use acquired resources, primarily C, N, and P from photosynthesis and root absorption for immediate structural growth (i.e., ‘acquisitive’) or divert a fraction of resources to storage structure reserves, such as starchy tubers, as a method of investing in survival at the expense of current growth (i.e., ‘conservative’; Chapin, 1980; Chapin et al., 1990). Annuals and perennials are typically characterized as having acquisitive and conservative strategies, respectively (Roumet et al., 2006). Consistent with this trend, relative growth rate, photosynthetic rate, nitrogen content, and specific leaf area are typically higher in annual than perennial species (Zangerl & Bazzaz, 1983; Garnier, 1992; Roumet et al., 1996; Poorter et al., 2012; Vico et al., 2016). Annuals have been further described as having higher reproductive effort, faster flowering and germination rates, and lower root mass fractions than perennials across various taxa (Gaines et al., 1974; Pitelka, 1977; Primack, 1979; Shipley & Parent, 1991; Milotic & Hoffman, 2016). Nevertheless, some annual and perennial species exhibit traits from both ends of the plant economics spectrum, which suggests more of a continuum than a dichotomy of trait combinations (Hancock & Pritts, 1987; Verboom et al., 2004; González-Paleo & Ravetta, 2015). Such trait variability is key in breeding new crops, and it is important for understanding how different annual and perennial wild populations respond to artificial selection for increased yield.

Artificial selection under cultivation can dramatically change annual and perennial trait variation. Cultivation, the act of planting and tending plants, often involves selective propagation of individuals with preferred features. Over time cultivated populations evolve in response to this artificial selection, leading to domestication: the evolution of morphological and genetic changes in cultivated populations relative to their wild progenitors (Harlan, 1995; Meyer et al., 2012). Domestication is a continuum, ranging from cultivated populations which display little or no differences relative to wild populations, to highly modified elite breeding lines, such as in contemporary maize (Harris, 1989; Bharucha & Pretty, 2010; Breseghello & Coelho, 2013). To determine the extent to which domestication has occurred, the precise identity of the wild ancestor of a crop and the exact cultivated populations derived from it are required. Because these data are not available for all species examined here, we use the term “cultivated” to refer to any accession that has a history of cultivation, acknowledging that this encompasses a broad spectrum of phenotypic and genetic change under artificial selection.

The “domestication syndrome,” a common suite of trait changes seen across different species in response to artificial selection for seed and/or fruit production (Hammer, 1984; Harlan, 1992), has been characterized for annual crops and woody perennial crops, but is generally lacking for herbaceous perennial species. In annuals, domestication typically includes loss of seed dormancy, higher germination, loss of shattering, greater seed size or number, and erect, determinate growth, among many others (Harlan et al., 1973; Olsen & Wendel, 2013; Abbo et al., 2014). In contrast to annuals, woody perennials often exhibit an extended juvenile phase, have an outcrossing mating system, and are usually clonally propagated; consequently, woody perennials have typically undergone fewer cycles of sexual selection under domestication and retain a greater proportion of wild genetic diversity relative to annual domesticates (Zohary & Spiegel-Roy, 1975; Miller & Gross, 2011; Gaut et al., 2015). Herbaceous perennial plants grown specifically for edible seeds were only recently targeted for selection (Suneson et al., 1963; Kantar et al., 2016, Crews and Cattani, 2018), with most breeding research occurring in the last few decades (DeHaan et al., 2016; Kane et al., 2016). Because relatively few herbaceous perennial species have been domesticated for seed production, it is unclear if they will follow an evolutionary trajectory similar to domesticated annual and woody perennial species, or if they will show a unique domestication syndrome.

Ongoing work seeks to understand how artificial selection for increased seed yield in herbaceous perennial species might impact vegetative traits and the capacity for perennation more generally. One hypothesis is that perennial seed crops are constrained by a vegetative-reproductive trade-off, where high reproductive allocation and sufficient storage allocation for perennation cannot coexist (Van Tassel et al., 2010). In other words, it may be possible for artificial selection to drive increases in seed yield in wild herbaceous perennial species, but those increases may cause losses in allocation to vegetative and perennating structures, resulting in a shift from perenniality to annuality (Denison, 2012; Smaje, 2015). An alternative hypothesis is that reproductive yield and vegetative biomass may be selected for in concert. The agricultural context provides an entirely new adaptive landscape that may allow novel combinations of traits in herbaceous perennial species (e.g., high reproductive output and longevity), combinations that might be unfavorable in natural environments (Crews & DeHaan, 2015; Cox et al., 2018).

The few studies of perennial seed crops undergoing artificial selection have demonstrated conflicting patterns of vegetative-reproductive covariation. In the perennial oilseed candidate *Physaria mendocina* (Brassicaceae), selection for seed yield was associated with lower root allocation and seed production in subsequent years compared to wild forms (González-Paleo et al., 2016; Pastor-Pastor et al., 2018). Similarly, in perennial wheat relatives (Poaceae), high seed production in year one was associated with lower yields and lower survival in subsequent years (Vico et al., 2016; Cattani, 2017). Nevertheless, concomitant perennation and high seed yield has been observed in multiple perennial cereals (Sacks et al., 2007; Jaikumar et al., 2012; Culman et al., 2013; Huang et al., 2018), suggesting that this combination is not biologically incompatible and may be specifically selected for in breeding.

Herbaceous perennial seed crop studies can benefit from juxtaposition with closely related annual domesticates, to clarify if domestication responses are dependent upon lifespan. Vico et al., (2016), in a meta-analysis of 67 annual-perennial pair studies across nine plant families, found greater reproductive allocation in annual crops and greater root allocation in perennial crops, which was the same pattern found in unselected wild groups. However, when absolute yield is considered, perennials produced an equivalent total seed weight to annuals, likely due to the perennial plants’ larger overall size (Vico et al., 2016). The perennial seed crops in this study were also just beginning to be domesticated in contrast to thousands of years of artificial selection in most annual crops. Such annual-perennial meta-analyses may be augmented by empirical research on specific lineages of plants, to determine if phylogenetically focused trends are similar to the broader patterns observed, as well as to allow a more precise biological interpretation. Lastly, a majority of the studies addressing congeneric lifespan trait differences are in the grass family (66%, as compiled by Vico et al., 2016); a more diverse phylogenetic sampling will be necessary to determine if observed patterns are ubiquitous.

Here we focus on the legume family (Fabaceae), which includes 19,500+ species of which more than 30% are predominantly herbaceous perennials (Ciotir et al., 2019). Fabaceae is the second most economically important plant family after the grasses (Poaceae), with 41 domesticated species dating back to the first agricultural systems, and 1,000+ species cultivated for various purposes across the world (Harlan, 1992; Lewis et al., 2005; Hammer & Khoshbakht, 2015). Legumes also account for at least one third of the world’s human dietary nitrogen and vegetable oil production (Vance et al., 2000; Graham & Vance, 2003). Fabaceae species therefore offer promising candidates for breeding and domestication of new perennial pulses (legume dry seed crops), but first, species of interest must be phenotypically characterized to narrow the selection process (Schlautman et al., 2018). To date, few herbaceous perennial legume species have been thoroughly assessed for agriculturally relevant traits such as seed size, germination rate, time to maturity, root/shoot allocation, and reproductive yield. Characterizing these and similar traits in herbaceous perennial crops and their wild relatives will be critical to assess what genetic variation is available for crop improvement through introgression and *de novo* domestication (Schlautman et al., 2018; Smýkal et al., 2018). *Lupinus* (lupine) and *Phaseolus* (common bean) were selected for this study based on the criteria that they include multiple annual and herbaceous perennial species, with availability of cultivated and wild seed accessions for both lifespans.

While there are known consistent differences in growth and allocation in some groups of annual and perennial species, gaps remain in our knowledge about how herbaceous perennials respond to artificial selection relative to their annual congeners in many plant lineages. Here we explore life history differences in *Lupinus* and *Phaseolus* by focusing on the following questions: 1) How do *wild* annual and perennial species allocate resources to seed production and vegetative growth? 2) What is the phenotypic signature of artificial selection in *cultivated* annual and perennial species? 3) What covariation exists between seed and adult vegetative growth traits, and is it consistent across lifespans and between cultivated and wild forms of a species? We address these questions by examining seed size, germination, and vegetative growth allocation among six annual and twelve perennial species of two economically important legume genera, *Lupinus* and *Phaseolus* (schematic of studied traits: **Figure 1**). Through this work, we hope to contribute to ongoing efforts characterizing life history differences in closely related annuals and perennials within Fabaceae, and shed light on how these lifespan groups may change with artificial selection.

**Figure 1.**
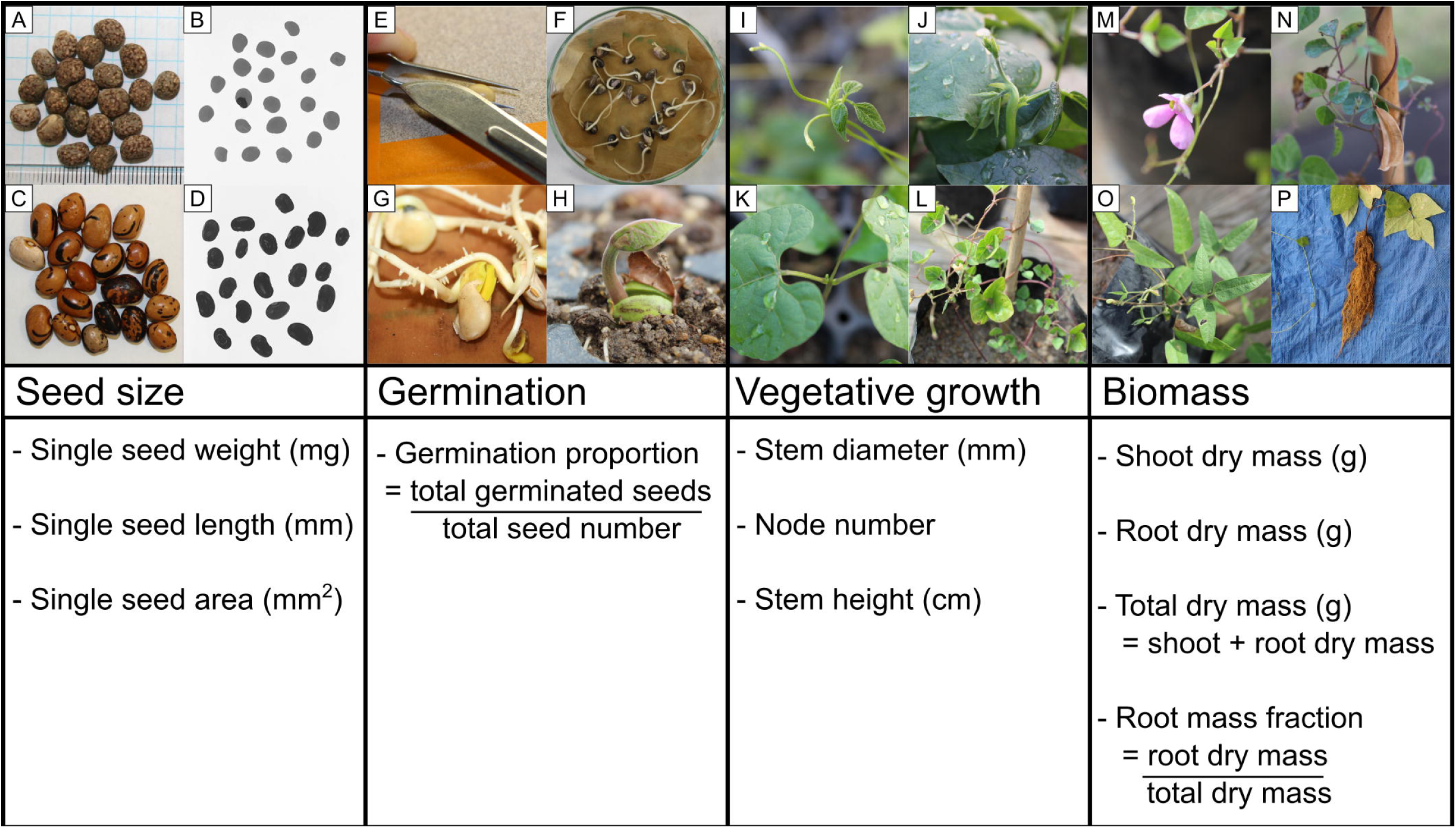
Schematic of the four developmental stages analyzed in this study: seeds, germination, early season vegetative growth, and late season biomass. **(A)** *Lupinus angustifolius* seeds; **(B)** *L. mutabilis* seeds prepared for analysis in ImageJ; **(C)** *Phaseolus coccineus* seeds; **(D)** *P. coccineus* seeds prepared for analysis in ImageJ; **(E)** scarification by nicking the seed coat; **(F)** *Phaseolus vulgaris* germinants; **(G)** *Lupinus albus* germinants; **(H)** *Phaseolus coccineus* germinant planted in greenhouse; **(I,J)** *Phaseolus* shoot apex at which stem height was measured to; **(K)** *Phaseolus* first node (unifoliate) below which stem diameter was measured; **(L)** fully grown *Phaseolus filiformis* shoot with developed nodes (unfolded leaves); **(M,N)** *P. filiformis* flower and ripe fruit; **(O)** *P. acutifolius* shoot biomass; **(P)** *P. coccineus* root biomass. Photo credit: SH.

## 2 MATERIALS AND METHODS

### 2.1 Plant material

For *Lupinus*, three annual species (*L. albus, L. angustifolius, L. arizonicus*; 35 accessions total) and eight perennial species (*L. albicaulis, L. albifrons, L. andersonii, L. elegans, L. mexicanus, L. mutabilis, L. polyphyllus, L. rivularis*; 28 accessions total) were included in the study. For *Phaseolus* three annual species (*P. acutifolius, P. filiformis, P. vulgaris*; 58 accessions) and four perennial species (*P. angustissimus, P. coccineus, P. dumosus, P. maculatus*; 74 accessions) were included (**Tables 1** & **2**). Species were chosen based on phylogenetic proximity and similar habitat types when possible; for *Lupinus* in particular, lifespan tends to segregate geographically and is generally phylogenetically disjunct (Drummond et al., 2012). Our sampling includes annual *Lupinus* species from the Mediterranean range (*L. albus, L. angustifolius*), with the remaining species from North and South America (Supplementary Table S5). Generally within *Lupinus*, annuality is the ancestral state, while perenniality evolved following oceanic dispersal to the Americas in response to novel montane environments (Drummond et al., 2012). *Phaseolus* species included here are more phylogenetically interspersed in terms of lifespan, with the perennials *P. coccineus* and *P. dumosus* and annuals *P. acutifolius* and *P. vulgaris* within the Vulgaris clade, the perennial *P. angustissimus* and annual *P. filiformis* within the Filiformis clade, and perennial *P. maculatus* alone in this study from the Polystachios clade (Delgado-Salinas, 2006). Geographically, our sampling of *Phaseolus* includes arid-adapted species of the Sonoran Desert (*P. acutifolius, P. angustissimus, P. filiformis*, and *P. maculatus*) and more tropically distributed species (primarily Mesoamerica: *P. coccineus, P. dumosus*, and *P. vulgaris*, with some; Supplementary Table S5).

**Table 1.**
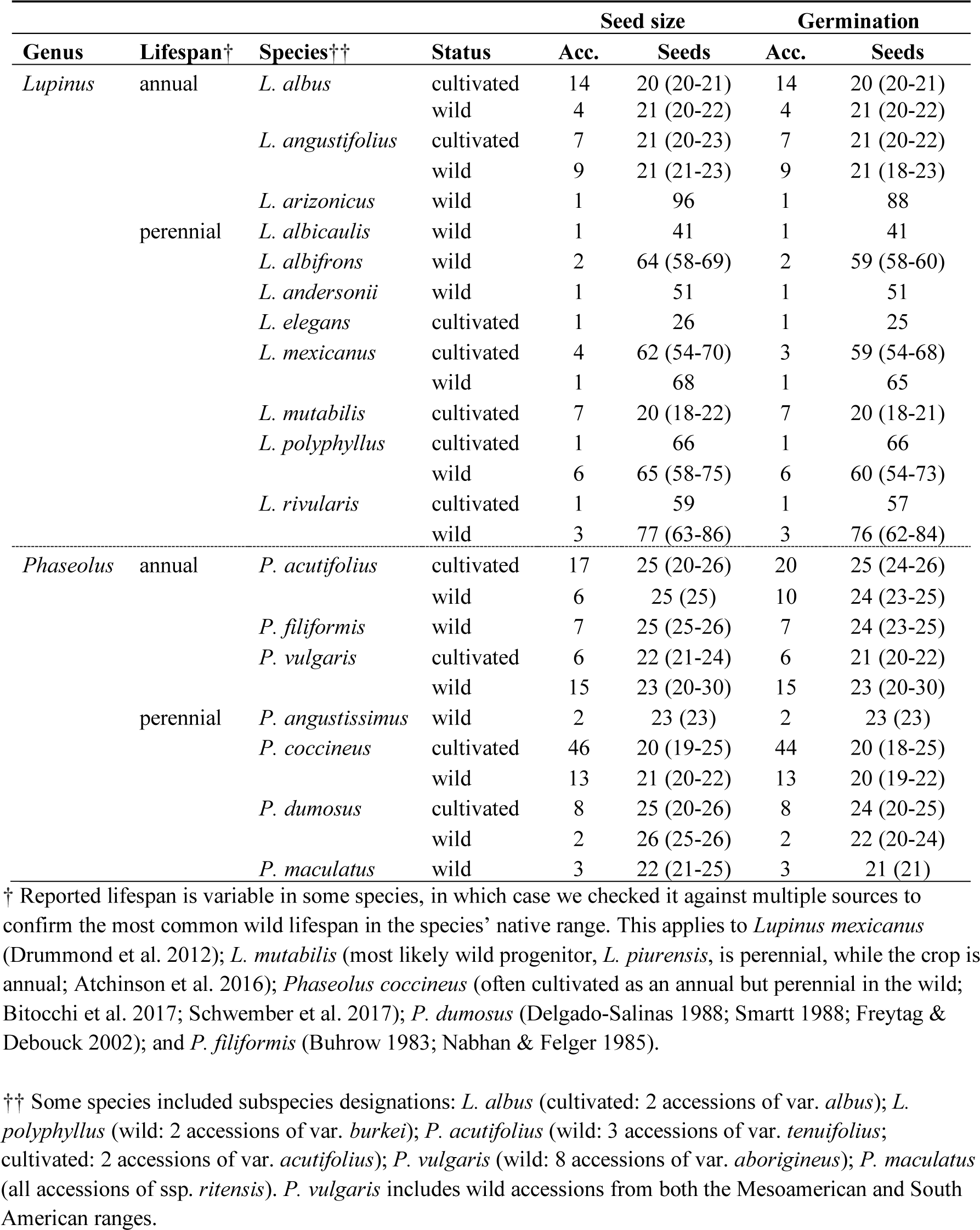
Summary of sampling for seed traits by species and cultivation status. ‘Acc.’ refers to the total number of accessions used for each species / cultivation status combination. ‘Seeds’ refers to the average seed number among accessions studied for seed size and germination (range of seed number in parentheses). Seed size collectively includes seed weight, length, and area.

In total 5,211 seeds (195 accessions) total were obtained from the United States Department of Agriculture’s National Plant Germplasm System (Western Regional PI Station, Pullman, WA, stored at −18°C) in spring 2016 and stored in a desiccator at 4°C, 33-50% relative humidity. All seeds were derived from plants regrown from the original collection material at the germplasm facility, but there are nevertheless possible genotype by environment and maternal effects that cannot be resolved here. 5,046 seeds were germinated and a subset of these grown from July to September 2016. The seed age for each accession, i.e., the length of time they were in frozen storage, ranged from one year to greater than 46 years, and a few accessions were of unknown age.

### 2.2 Lifespan and cultivation status assignment

Species lifespan data were obtained from USDA accession descriptions and double-checked against the literature. In rare cases where USDA information contradicted generally accepted botanical knowledge, the selected lifespan was verified using multiple species databases and publications (see notes in **Table 1**). We classified species in terms of the predominant lifespan in the wild in its native range. Cultivation status was taken directly from the USDA’s description, with the descriptors ‘cultivated’, ‘cultivar’, and ‘landrace’ all categorized as ‘cultivated’ for the purpose of this dataset. We use the umbrella term ‘cultivated’ rather than ‘domesticated,’ since we do not have the data to determine the extent to which phenotypic and genetic change has occurred from the original wild population selected upon (see Introduction). Cultivated annual *Lupinus* included here originated in the Mediterranean region, while cultivated perennial *Lupinus* originated in the Americas (Wolko et al., 2011). All *Phaseolus* species are native to the Americas and were originally cultivated in either Mesoamerica or South America with some later selection occurring in Eurasia (Bitocchi et al., 2017). Our dataset includes cultivated accessions from both regions but primarily the American region, as well as four of the five domesticates in the genus. Cultivated species here were domesticated at least 1000 years before present (BP), most much longer, except for several *Lupinus* species: *L. angustifolius* (200 years BP) and *L. elegans, L. mexicanus, L. rivularis*, and *L. polyphyllus* (less than 100 years BP). Species-specific details on geographic origin and domestication are available in Supplementary Table S5.

### 2.3 Traits measured

#### 2.3.1 Seed size

A total of 2,034 annual seeds (86 accessions) and 3,005 perennial seeds (102 accessions) were analyzed for size traits (**Figure. 1**; **Table 1**). Seeds from each accession were weighed in bulk to the nearest mg and mean single seed weight was estimated by dividing the bulk weight by the total number of seeds for that accession. We imaged all accessions on a light table with a fixed camera at a resolution of 640×360 or 1349×748 pixels (differences accounted for in linear models). We used ImageJ (Rasband, 1997-2016) to measure mean single seed length and area.

#### 2.3.2 Germination

2,192 annual seeds (93 accessions) and 2,854 perennial seeds (99 accessions) were monitored for germination (number discrepancy is due to a few germinated accessions not being analyzed for seed size and vice versa, and some seed loss in the germination procedure). Number of germinated seeds was monitored for each accession over time and used to calculate germination proportion (**Figure 1**; **Table 1**). Seeds were germinated on RO-water dampened germination paper or with a dampened cotton ball in petri dishes, following 1) sterilization by soaking in 1% bleach for two minutes and rinsing, 2) scarification by nicking the seed coat on the flat side the seed with a scalpel (to break physical dormancy, i.e., a water-impermeable seed coat), and 3) soaking for an average of 23 hours (range: 9-33 hours) by submersion in RO water, kept at 21-24°C. The soaking start time was considered time point 0 (i.e., the sowing date) for germination counts, since all necessary resources were available for seeds to germinate. Some *Lupinus* seeds were unsterilized, germinated on paper in trays, and, rarely, scarified with P100 to P150 sandpaper; this variation was accounted for in downstream analyses. The germination apparatus was placed onto a 24°C heat mat in 24-hour dark conditions (except for germinant counts; López Herrera et al., 2001). Germination was defined as an extension of the radicle past the seed coat. In rare cases where the seed coat was lost or very hard, it was defined as a distinct vigorous movement of the radicle away from the seed or a distinct pushing of the seed coat away from the seed, respectively. Seeds were treated for fungal infection with tea tree oil or Banrot 40WP fungicide solution (prepared according to the product label). Infected, potentially salvageable seeds were soaked in 1-2% bleach and rinsed with RO water. Germination paper was re-moistened with RO-water when dry (approximately every two days). Germinated seeds were also scored for seed quality following any potential damage from pre-germination treatments (0-2, with 2 being the highest quality), and the number of seeds which were compromised due to procedural damage or damage in storage (removed from analysis). Any variation in germination protocol was noted and accounted for in statistical models, as well as the covariates seed quality and seed age (duration of frozen storage at the germplasm center since the last seed increase). Germinated seed number was observed until all seeds had germinated or died, up to 27 days. Subsets of individuals from the original number germinated were chosen for further growth measurement based on the presence of ≥10 vigorous individuals, if the accession was from the native range of the species (preferred), and if the accession was from a duplicate geographic location (removed). Due to the low sample size of species and accessions with viable individuals, *Lupinus* was excluded for all post-germination trait analyses in this study.

#### 2.3.3 Vegetative growth measurements

507 annual individuals (43 accessions) and 232 perennial individuals (29 accessions) of *Phaseolus* were transplanted to a greenhouse and measured for at least one vegetative trait three weeks after transplanting (details below; **Figure 1**; **Table 2**). Seedlings were initially planted on July 13-15, 2016 in a mixture of unsterilized local riverine soil (Smoky Hill River, Salina, KS: 38.765948 N, -97.574213 W) and potting soil (PRO-MIX) in small trays until they could be planted in 8” tall x 4” wide bag pots after 1-2 weeks. Initial planting date in small trays was used as the baseline for future measurements (days after planting, DAP), since despite different sowing dates, they were developmentally similar upon planting. Bag pots were filled with a mixture of the same riverine soil and coarse sand, to mimic field soil while also maximizing drainage. All *Phaseolus* were twining and were trained up four-foot bamboo poles. Plants did not receive any rhizobial inoculant treatment. Plants were initially bottom-watered twice daily (10AM, 6PM) for 20 minutes. On August 18, 2016, watering was changed to once for 10 minutes every two days. A shade cloth was incorporated in the greenhouse until August 23, 2016. 90 of the most vigorous *Phaseolus* plants were moved from the greenhouse to the outdoors on August 25-26, 2016 to expose them to a more light-intense, natural environment. At this time, individual plants in both the greenhouse and outdoors were randomized to reduce spatial bias.

**Table 2.**
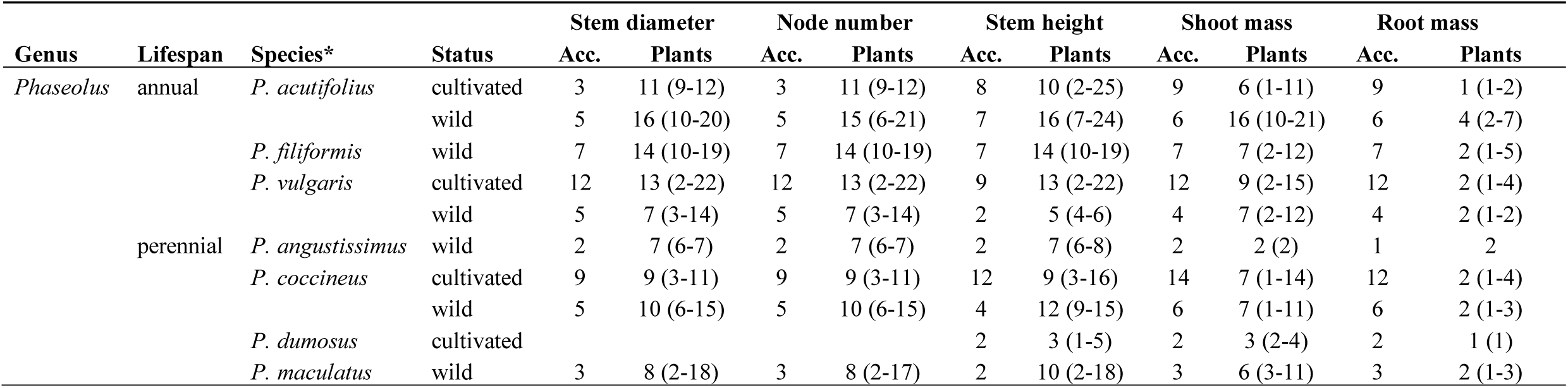
Summary of sampling for vegetative growth and biomass traits for *Phaseolus*, by species and cultivation status. “Acc.” refers to seed accession count. “Plants” refers to the average plant number among accessions studied for that trait (range of plant number in parentheses).

All growth analyses were conducted at the research facilities of The Land Institute (Salina, KS). At 19-23 DAP (25-40 days after sowing), plants were measured for stem diameter below the first node, total developed node number counted from the unifoliate node to the last node with an unfolded leaf, and stem height from ground to shoot apex on the tallest main stem, with twining stems being uncoiled from their poles and straightened as far as possible without damaging the plant. Plants were checked for reproductive status before biomass harvest. At 68-75 DAP (74-93 days after sowing), a random subset of plants was harvested for shoot and root (washed) biomass (**Figure 1; Table 2**), which was dried at a minimum of 37°C for at least 24 hours. Biomass was weighed on a precision or analytical scale depending on plant size. Root mass fraction was calculated on an individual plant basis from biomass measurements (root dry mass / total dry mass). Variation in growth conditions (greenhouse or outdoors), plant health (ordinal rating: 0,1,2; unhealthy, moderate health, healthy), and reproductive status (ordinal rating: 0,1,2,3; no reproduction, budding, flowering, fruiting) were recorded for individual plants and accounted for in statistical analyses (see below).

### 2.4 Statistical analyses

In order to assess associations between lifespan, cultivation status, and trait variation, we used linear models and *post-hoc* comparative analyses on a set of mean values for each trait from each accession (**Tables 3, 4**). Associations of lifespan, genus, cultivation (nested within genus and lifespan), and species (nested within genus, lifespan, and cultivation), in addition to any relevant covariates for the focal trait, were tested using linear models. All potentially confounding factors were included in the original model and were then dropped sequentially if found to be nonsignificant. The base model for all main analyses was: *trait = genus* + *lifespan* + *genus*\**lifespan* + *genus/lifespan/cultivation* + *genus/lifespan/cultivation/species*. Analyses were calculated for mean accession-level data for all traits and covariates. Seed and germination trait analyses were conducted for *Lupinus* and *Phaseolus*, but vegetative growth analyses included only *Phaseolus* (see Growth measurements section above). Due to concerns about lifespan lability of *P. dumosus*, linear models for all traits were checked with this species included and then with the species removed from the dataset. Pairwise comparisons of lifespan and cultivation effects for each trait were evaluated with *post-hoc* Tukey HSD tests (Supplementary Table S1).

**Table 3.** Results of linear models for seed traits for a combined *Lupinus* and *Phaseolus* dataset. Letters denote separate models with different covariates, while the main effects are the same for all traits: (a) seed size traits and (b) germination.

|  | Trait | Lifespan | Genus | Lifespan ×<br>Genus | Cultivation | Species |  |  |  |  |
| --- | --- | --- | --- | --- | --- | --- | --- | --- | --- | --- |
| (a) | Seed weight | $F_1 = 55.27^{***}$ | $F_1 = 33.52^{***}$ | $F_1 = 93.73^{***}$ | $F_4 = 34.80^{***}$ | $F_{19} = 1.16$ | | | | |
| | Seed length | $F_1 = 71.05^{***}$ | $F_1 = 39.12^{***}$ | $F_1 = 200.80^{***}$ | $F_4 = 60.85^{***}$ | $F_{18} = 5.13^{***}$ | | | | |
| | Seed area | $F_1 = 63.91^{***}$ | $F_1 = 12.58^{***}$ | $F_1 = 133.33^{***}$ | $F_4 = 38.66^{***}$ | $F_{19} = 2.16^{**}$ | | | | |
|  | Trait | Lifespan | Genus | Lifespan ×<br>Genus | Cultivation | Species | Age | Scarification | Soak time | Seed quality |
| (b) | Germination<br>proportion | $F_1 = 36.30^{***}$ | $F_1 = 2.69$ | $F_1 = 11.88^{***}$ | $F_4 = 3.67^{**}$ | $F_{18} = 2.27^{**}$ | $F_1 = 5.72^*$ | $F_1 = 0.39$ | $F_1 = 1.56$ | $F_1 = 4.66^*$ |

\* $P < 0.05$ ; \*\* $P < 0.01$ ; \*\*\* $P < 0.001$ .

**Table 4.**
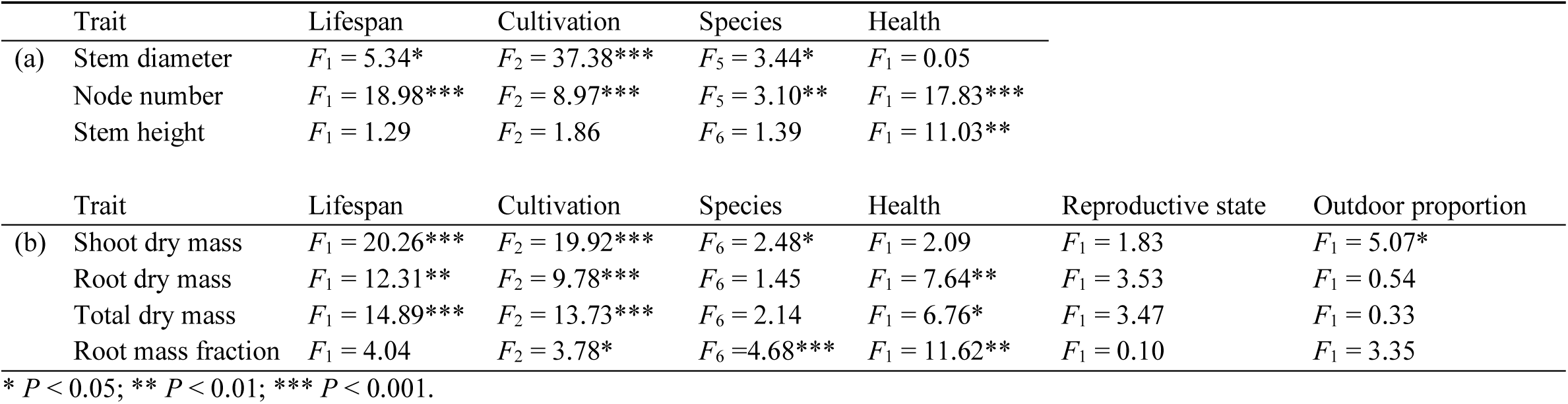
Results of linear models for vegetative growth traits in *Phaseolus*. Letters denote separate models with different covariates, while the main effects are the same for all traits: (a) early vegetative growth traits and (b) biomass traits.

| Trait | Lifespan | Cultivation | Species | Health |  |  |
| --- | --- | --- | --- | --- | --- | --- |
| (a) Stem diameter | $F_1 = 5.34^*$ | $F_2 = 37.38^{***}$ | $F_5 = 3.44^*$ | $F_1 = 0.05$ | | |
| Node number | $F_1 = 18.98^{***}$ | $F_2 = 8.97^{***}$ | $F_5 = 3.10^{**}$ | $F_1 = 17.83^{***}$ | | |
| Stem height | $F_1 = 1.29$ | $F_2 = 1.86$ | $F_6 = 1.39$ | $F_1 = 11.03^{**}$ | | |
| Trait | Lifespan | Cultivation | Species | Health | Reproductive state | Outdoor proportion |
| (b) Shoot dry mass | $F_1 = 20.26^{***}$ | $F_2 = 19.92^{***}$ | $F_6 = 2.48^*$ | $F_1 = 2.09$ | $F_1 = 1.83$ | $F_1 = 5.07^*$ |
| Root dry mass | $F_1 = 12.31^{**}$ | $F_2 = 9.78^{***}$ | $F_6 = 1.45$ | $F_1 = 7.64^{**}$ | $F_1 = 3.53$ | $F_1 = 0.54$ |
| Total dry mass | $F_1 = 14.89^{***}$ | $F_2 = 13.73^{***}$ | $F_6 = 2.14$ | $F_1 = 6.76^*$ | $F_1 = 3.47$ | $F_1 = 0.33$ |
| Root mass fraction | $F_1 = 4.04$ | $F_2 = 3.78^*$ | $F_6 = 4.68^{***}$ | $F_1 = 11.62^{**}$ | $F_1 = 0.10$ | $F_1 = 3.35$ |

\* $P < 0.05$ ; \*\* $P < 0.01$ ; \*\*\* $P < 0.001$ .

To assess trait covariation within the whole dataset, Pearson product-moment correlations were performed on all pairwise combinations of the 11 measured traits using mean accession-level data, to determine the magnitude and direction of their relationships, excluding *Lupinus* for post-germination traits. Pearson correlations were also run on subsets of the data to qualitatively assess any trait covariation differences for the following paired groups: annual vs. perennial, cultivated vs. wild, and *Lupinus* vs. *Phaseolus* (Supplementary Figure S1). Each subset of the data was only restrictive in regard to one criterion, e.g., the annual subset contained data for both cultivated and wild, *Lupinus* and *Phaseolus*, annual accessions. Lastly, we ran linear models and Tukey HSD tests for wild accessions of each genus to determine the impact of broad geographic/habitat differences in species distributions: for *Lupinus*, these were Mediterranean annual vs. American perennial (and one annual, *L. arizonicus*, which had to be excluded from Tukey tests); for *Phaseolus*, these were desert vs. tropical (Supplementary Tables S2, S3, S4). The base model for the geographic analyses was: *trait = geography* + *lifespan* + *geography*×*lifespan* + *geography/lifespan/species*, including any relevant covariates. See Supplementary Table S5 for the assignment of distribution categories to species. All statistical analyses were performed in R v. 3.6.1 (R Core Team, 2019).

## 3 RESULTS

We investigated annual and perennial species in *Lupinus* and *Phaseolus* for potential lifespan differences and covariation in seed size, germination, and vegetative growth. Our overall finding was that in both *Lupinus* and *Phaseolus*, annual and perennial species exhibited differences for most seed traits. Further, cultivated accessions had greater trait values compared to wild accessions for almost all traits measured, regardless of genus and lifespan. Lastly, most traits examined here are positively correlated, regardless of genus, lifespan, and cultivation status.

### 3.1 Trait differences in wild annual vs. perennial *Lupinus* and *Phaseolus* accessions

Wild annual *Lupinus* species had significantly larger seeds and significantly higher germination proportion than wild perennial *Lupinus* species (**Figures 2A-D**; see Supplementary Table S1 for all mean values, standard deviation, and Tukey test significance). Wild annual *Phaseolus* species had smaller seeds than wild perennial *Phaseolus* species (nonsignificant), but like *Lupinus*, wild annual germination proportion was significantly higher than that of wild perennial *Phaseolus* (**Figures 2A-D**). Although nonsignificant, wild perennial *Phaseolus* mean seed weight (101 mg) was nearly twice that of wild annuals (59 mg; **Figure 2A**; Supplementary Table S1). Wild annual *Phaseolus* had similar to nonsignificantly larger vegetative trait values compared to wild perennials, with the largest relative differences seen in stem height (31.94 cm annual vs. 22.38 cm perennial) and total dry mass (1.48 g annual vs. 0.99 perennial; **Figures 2G, J**; Supplementary Table S1). Mean root mass fraction was equivalent for wild annual and perennial *Phaseolus* (0.22; **Figure 2K**; Supplementary Table S1).

**Figure 2.**
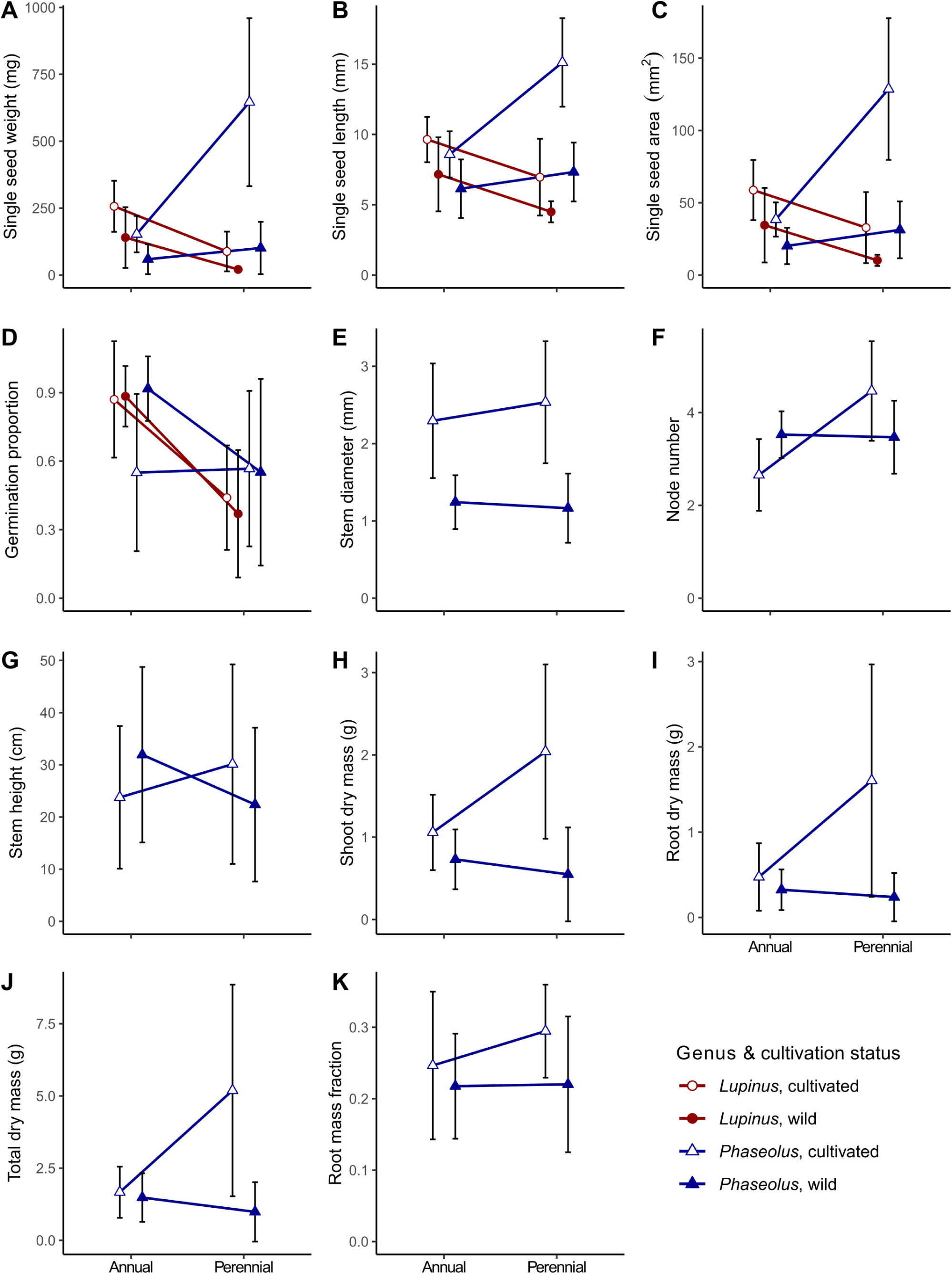
Panel of differences among lifespan, genus, and cultivation status for the 11 focal traits: **(A-C)** seed size traits; **(D)** germination proportion; **(E-G)** early vegetative growth traits; **(H-K)** shoot and root biomass traits. Central points represent means of all accessions for that category; error bars represent one standard deviation (absence of error bars for wild perennial *Lupinus* in **(A)** is due to relatively very small standard deviation).

While our data do not allow precise interpretation of geographic effects, linear models including geographic location as a main effect along with lifespan (for only wild accessions) found that geography explained a significant portion of the variation seen in all seed size traits for wild accessions in both genera (Supplementary Table S2). In *Phaseolus* geography was significant for germination proportion, stem diameter, stem height, shoot dry mass, root dry mass, total dry mass, and root mass fraction (Supplementary Tables S2, S3). *Post-hoc* Tukey tests indicated that Mediterranean annual *Lupinus* showed significantly larger seed size (all traits) and significantly higher germination proportion compared to American perennial *Lupinus*. Tropically distributed *Phaseolus* accessions had larger seeds, stem diameter, stem height, and dry biomass than desert-distributed species; compared to desert perennials, desert annuals more often had significantly smaller traits than tropical *Phaseolus* species; Supplementary Table S4). Within both tropical and desert *Phaseolus* species, perennials tended to have greater seed size than annuals, but generally smaller or similar vegetative trait sizes (Supplementary Table S4). Mean germination proportion was relatively high (> 0.85) for all *Phaseolus* groups except tropical perennials (0.40, although tropical perennials also had the greatest standard deviation; Supplementary Table S4).

Overall, wild annual and perennial *Lupinus* and *Phaseolus* species show contrasting lifespan differences in seed size but have similar germination patterns. These patterns were supported by a strongly significant lifespan×genus interaction for all seed traits in our linear model (**Table 3**). In the complete dataset, lifespan alone explained a significant amount of the variation seen in all traits with the exception of stem height and root mass fraction (**Tables 3**, **4**). In our linear models, scarification and soak time were not significant predictors of germination; age and seed quality had significance at *P* < 0.05. Plant health was significant in the linear models for all vegetative and biomass traits except stem diameter. Reproductive state and outdoor proportion were not statistically significant in the linear models for any biomass traits, except in the case of shoot dry mass, where outdoor proportion was significant at *P* < 0.05. The removal of *P. dumosus* from the dataset (see Methods) changed some linear model results for biomass traits. For shoot dry mass, the species effect went from significant at *P* < 0.05 to nonsignificant. For root and total dry mass, reproductive state went from nonsignificant to marginally significant at *P* < 0.1. For root mass fraction, the cultivation effect went from significant at *P* < 0.05 to nonsignificant, and outdoor proportion went from nonsignificant to marginally significant at *P* < 0.1.

### 3.2 Trait differences in cultivated vs. wild *Lupinus* and *Phaseolus* accessions

Cultivated annual and cultivated perennial *Lupinus* species had larger seeds than their wild relatives. There was no significant change in germination with cultivation for either annual or perennial *Lupinus* (**Figures 2A-D;** Supplementary Table S1). Similarly, cultivated annual and perennial *Phaseolus* species had larger seeds compared to wild relatives (**Figures 2A-K**). Cultivation differences in seed size were nonsignificant for annual *Phaseolus*, although cultivated annual seed weight (153 mg) was nearly three times larger than wild annuals (59 mg) (**Figure 2A**; Supplementary Table S1). Cultivated perennial *Phaseolus* had significantly greater seed size in all traits with seed weight over six times larger in cultivated perennials (646 mg) than wild perennials (101 mg) (**Figure 2A**; Supplementary Table S1). Germination proportion was significantly lower in cultivated annual *Phaseolus* (0.55) relative to wild annuals (0.92), while perennial germination remained at approximately the same low proportion regardless of cultivation status (0.57 cultivated vs. 0.55 wild; **Figure 2D**; Supplementary Table S1).

In addition to larger seeds and steady or reduced germination rates, cultivated *Phaseolus* species exhibited slight to large changes in vegetative features as well, i.e., stem diameter, node number, and stem height, as well as shoot dry mass, root dry mass, and their derivatives (**Figures 2E-K**). Cultivated perennial *Phaseolus* species tended to have considerably larger vegetative features relative to their wild relatives, whereas annual *Phaseolus* species displayed more subtle cultivation differences in vegetative features compared to their wild relatives (**Figures 2E-K**; Supplementary <u>Table S1</u>). Stem diameter was significantly greater in both annual and perennial *Phaseolus* (**Figure 2E**; Supplementary Table S1). Cultivated annual *Phaseolus* showed nonsignificantly lower values in node number and stem height compared to wild annuals, while cultivated perennial *Phaseolus* showed nonsignificantly larger values in both traits compared to wild perennials (**Figures 2F,G**; Supplementary <u>Table S1</u>). Both cultivated annual and perennial *Phaseolus* had greater dry biomass trait values than their wild relatives. For cultivated annual *Phaseolus*, the only significantly larger biomass value was shoot dry mass (1.06 g cultivated vs. 0.73 g wild; **Figure 2H**; Supplementary <u>Table S1</u>). In contrast, all biomass trait values were significantly greater in cultivated relative to wild perennial *Phaseolus* (except root mass fraction), with a greater than six-fold larger root dry mass value (1.60 g cultivated vs. 0.24 g wild; Figure 2**I;** Supplementary <u>Table S1</u>).

In summary, cultivated annual and cultivated perennial accessions of *Lupinus* and *Phaseolus* showed greater seed size, similar germination, and greater vegetative size (for *Phaseolus*) compared to their wild counterparts. In our linear model, cultivation likewise explained a significant amount of variation for all traits except stem height (**Tables 3, 4**).

### 3.3 Phenotypic covariation across all traits

Most traits measured in this study showed positive correlations within the total dataset (**Figure 3**) and each subset of the data (Supplementary Figures S1A-F). Considering the entire dataset, all seed dimensions (weight, length, area) were nearly perfectly correlated (R^2^ = 0.95-0.98; **Figure 3**). Seed traits were also significantly positively correlated with all vegetative growth traits, having the highest correlation with stem diameter and most biomass characters (R^2^ = 0.65-0.79; **Figure 3**). Stem diameter had positive albeit non-significant correlations with node number and stem height, and significant positive correlations with all biomass traits. Node number and stem height had nonsignificant positive correlations with most biomass traits (**Figure 3**). Biomass traits including shoot mass, root mass, total mass, and root mass fraction were significantly correlated with one another, and there was a tight relationship between shoot and root dry mass (R^2^ = 0.84; **Figure 3**). Germination proportion was the only trait in the total dataset to have no significant correlations and to have negative correlations with other traits (**Figure 3**).

**Figure 3.**
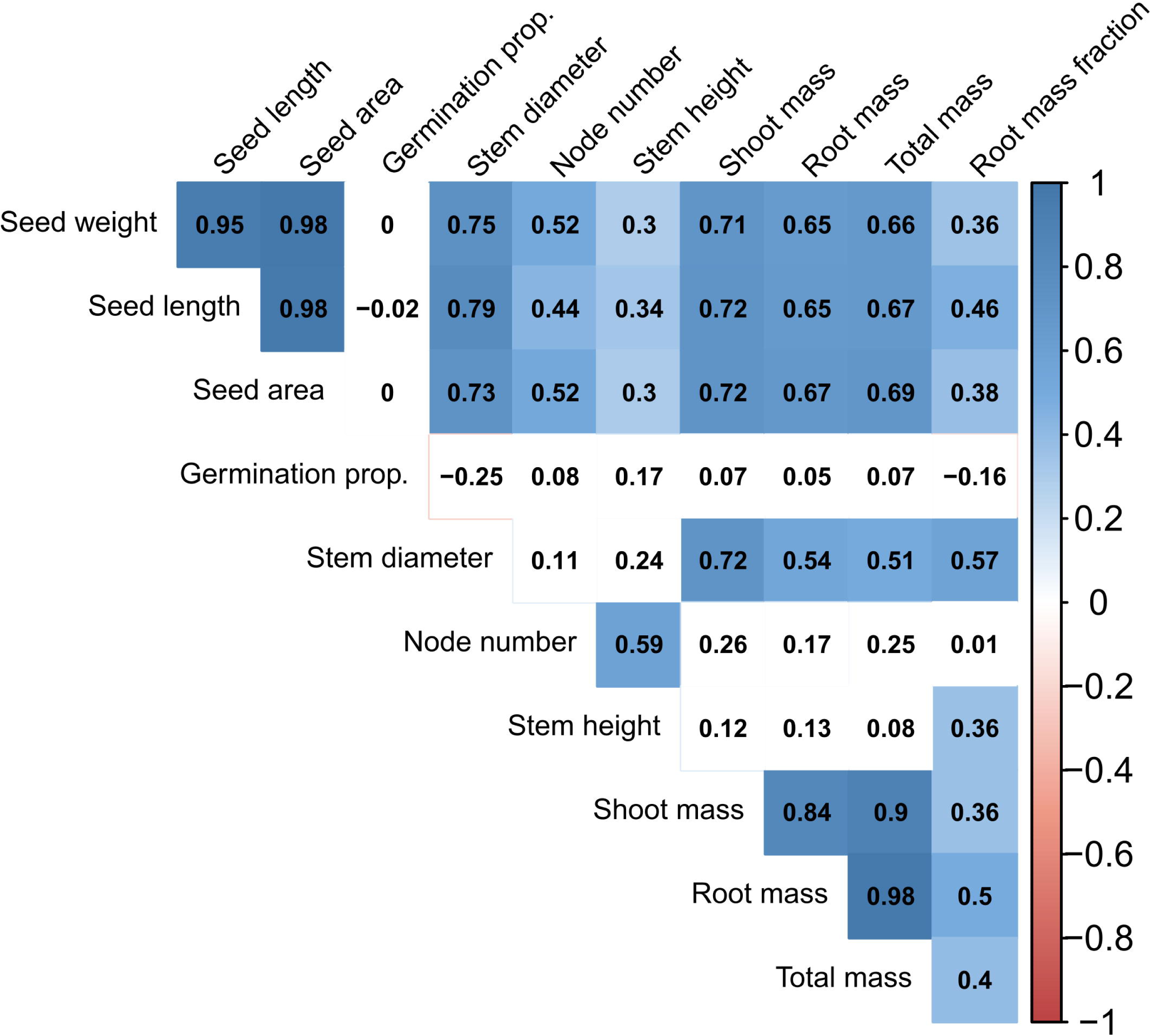
Correlation diagram for all traits for all accessions in the dataset. Seed traits and germination include both *Lupinus* and *Phaseolus*, whereas vegetative traits (stem diameter to root mass fraction) include only *Phaseolus* data. Numbers in boxes represent the Pearson correlation coefficient. Blue and red colors indicate significant positive and negative correlations (at the α = 0.05 level), respectively; absence of color signifies lack of significance.

Subsets of our data also displayed similarly positive trait correlations with some exceptions. Due to sample size, each subset was restricted in regard to only one criterion (genus, lifespan, or cultivation status), e.g., the perennial subset contained both cultivated and wild, *Lupinus* and *Phaseolus*, perennial accessions. Notably, the annual subset (including both cultivated and wild annuals of *Lupinus* and *Phaseolus*) showed negative correlations between node number and most traits (significant for seed size traits, stem diameter, and shoot mass), as well as a significant negative correlation between germination proportion and stem diameter (Supplementary Figure S1A). In contrast, the perennial subset, the wild subset, and cultivated subset respectively tended to show positive correlations for the trait pairings which were negatively correlated in annuals (Supplementary Figure S1B-D). Significant positive seed size to vegetative trait correlations were stronger and more common in the perennial and cultivated subsets than the annual and wild subsets (Supplementary Figures S1B,D). In the *Lupinus* subset, all seed size traits were significantly positively correlated with germination proportion, whereas in the *Phaseolus* subset, all seed size traits had nonsignificant negative correlations with germination proportion (Supplementary Figures S1E,F).

Overall, trait correlations for the total dataset and each subset were predominantly positive, most consistently among seed size traits and among vegetative traits, with most negative correlations occurring for germination proportion and node number.

## 4 DISCUSSION

This study examined lifespan and cultivation effects among annual and herbaceous perennial legume species in two genera, with the aim of identifying common trends and potential covariation among traits. Consistent trends included higher germination proportion in annual than perennial accessions, greater seed and plant size in cultivated relative to wild accessions for both annual and perennial species, and positive correlations among most seed and vegetative traits for annual and perennial species, both in the wild and under cultivation.

### 4.1 Wild annuals and perennials differ in seed size and germination proportion

Wild annual *Lupinus* species had larger seeds than wild perennial *Lupinus* species in our dataset. Larger seed size in wild annual *Lupinus* may be explained by divergent evolutionary histories, as the largest wild seeds were of our annual Mediterranean species (*L. albus* and *L. angustifolius*) in contrast to smaller-seeded perennial species distributed across the Americas (Supplementary Tables S4, S5). Such geographic differences are likely due to fine-scale genetic and environmental factors which shaped the evolutionary path of each species. Annual *L. albus*, for instance, likely has large seeds due to its distinctly early flowering time, which allows for a longer reproductive phase and thus larger pods and seeds (Berger et al., 2017). Considering North America alone, in a different set of three western *Lupinus* species, seed size was found to be higher in the herbaceous perennial species than the annual species, although the perennial had fewer seeds per pod and lower reproductive effort (Pitelka, 1977). This suggests that seed size is not necessarily proportional to total reproductive allocation but is instead impacted by other forms of selection, such as early competition for light and nutrients (Thompson & Hodkinson, 1998). Seed weight has also been seen to vary by five-times in a single population of annual *Lupinus texensis* (Schaal, 1980), and, in perennial species, seed size can vary substantially throughout the lifespan of an individual plant as well (e.g., *L. polyphyllus*; Aniszewski et al., 2001). Thus, the seed size variation we observed must also be considered in the context of the original maternal plant sampled and multiple intra-individual factors including age, pod position on the plant, and seed size-number trade-offs, as well as environmental factors such as soil fertility and climate. While lifespan was a significant predictor of seed size in our linear models, lifespan-dependent seed size patterns in *Lupinus* will benefit from further investigation in environmentally specific contexts, especially more closely related annual and perennial species in North and South America.

Data presented here demonstrated higher germination proportion in wild annual *Lupinus* relative to wild perennial congeners. Similarly high germination has been observed in other studies of annual *Lupinus* species (Schaal, 1980; including in *L. albus* after 26 years in frozen storage, Dobiesz & Piotrowicz-Cieślak, 2017), and similarly low germination proportion has been observed in other perennial western North American *Lupinus* species (Greipsson & El-Mayas, 2003; Sõber & Ramula, 2013). Pitelka (1977) found that while North American herbaceous perennials had somewhat higher germination than annual *Lupinus* in field sites, both groups had less than 40% germination. Our germination differences could also be impacted by the lower mean age (duration of frozen storage at the germplasm center) of wild annual (9 years) compared to wild perennial *Lupinus* (23 years) accessions, although cultivated *Lupinus* accessions showed the same germination pattern with a smaller storage age difference (7 years annual vs. 11 years perennial). Seed age also did not explain a large amount of the germination variation (**Table 3**). It is possible that some germination requirements were unmet for perennial *Lupinus*; however, any cold requirements were likely fulfilled given extended periods in frozen storage, and scarification successfully breached the seed coat in every accession.

In contrast to *Lupinus*, the wild perennial *Phaseolus* species in this study exhibited greater seed size trait values relative to annual *Phaseolus* species, which is consistent with the general life history expectation that later successional species produce larger seeds that are better able to compete for resources (Thompson & Hodkinson, 1998). This trend was consistent for *Phaseolus* species native to the desert and the tropics (Supplementary Table S4). Tropical *Phaseolus* species had higher mean seed size than desert *Phaseolus*, which is consistent with the broad pattern of increasing seed size at lower latitudes (Moles et al., 2007; Supplementary Table S4). The large range of phenotypic variation observed in annual and perennial *Phaseolus* (**Figures 2A-K**) is also consistent with the large distributions of several species, such as *P. vulgaris* and *P. coccineus*, which inhabit environments ranging from very arid to very humid (Dohle et al., 2019).

While few studies have compared germination in wild *Phaseolus* species, Bayuelo-Jiménez et al. (2002) reported a similarly high germination proportion (0.85+) for scarified wild annual and perennial *Phaseolus* accessions, including the desert perennials *P. angustissimus* and *P. filiformis*. Even though annuals showed higher mean germination than perennials in this study, desert perennial accessions all reached 100% germination, suggesting that the broader trend is more applicable to the tropical species studied here (Supplementary Table S4). Our lowest wild germination was observed in the tropical perennials *P. coccineus* and *P. dumosus* (Supplementary Table S4), which is consistent with the trend that seed dormancy is less common in tropical herbaceous species in regions of higher rainfall (Baskin & Baskin, 2014).

While germination showed consistent differences across genera, we cannot make ecologically robust conclusions here, since we effectively removed the legumes’ primary form of physical dormancy through scarification, and each accession was stored frozen for different lengths of time at USDA facilities, which can have diverse effects on species’ germination biology (Walters et al., 2005). Our results are likely more reflective of the viability of seeds after long-term storage. In this light, our findings are consistent with life history predictions that annual species will maintain a complex, long-lived seed bank in which dormancy breaking and germination is staggered over time to ensure offspring survival in variable environments (Venable & Lawlor, 1980; Gremer et al., 2016). Perennials may be less selectively constrained by this pressure due to the parent plant’s persistent survival in more stable environments (Cohen, 1966; Thompson et al., 1998). Physical dormancy in legumes is reinforced by increased impermeability of the seed coat and drying of the seed, although the physiological cost of increased seed dormancy and chemical inhibitors are unresolved for most of the species included here (Baskin & Baskin, 2014).

The lack of significant differences in vegetative traits between wild annual and perennial *Phaseolus* suggests that the traits studied here do not diverge at this growth stage among different *Phaseolus* life history strategies in nature. Perennials’ somewhat lower mean vegetative growth could be due to a slower growth rate and the early stage of growth at which the traits were measured (three weeks after planting), since some perennials have been shown to be able to achieve a higher total biomass than related annuals when the entire growing season is considered (Dohleman & Long, 2009). Previous studies have also found that vegetative growth, root, and shoot biomass are similar between closely related annuals and perennials, up to 40 days of growth, before their resource allocation patterns diverge (De Souza et al., 1987; Garnier, 1992). The significance of plant health in vegetative linear models for stem height and node number may reflect a greater sensitivity of response in stem length traits to environmental stressors compared to stem diameter; the health effect on root dry mass and derived traits (total dry mass and root mass fraction) may be due to a lower sample size for these traits compared to shoot dry mass alone (**Tables 2, 4**).

Wild *Lupinus* and *Phaseolus* differed in their lifespan-associated seed size patterns but were similar in germination patterns. Inconsistent lifespan-associated differences may be explained by the genera’s separate evolutionary histories: *Lupinus* and *Phaseolus* are in distant taxonomic tribes, few of the species in this study have overlapping geographic distributions, and phylogenetic clustering of their annual and perennial species is different. Per the last point, annuals and perennials clustered in distant clades within a genus may be phenotypically different due to a different common ancestor with a separate evolutionary history, while differences between annuals and perennials interspersed within the same clade are more likely be intrinsically lifespan-related (Vico et al., 2016). In our study, annuals and perennials in *Lupinus* were typically in separate clades (consistent more generally across the genus; Drummond et al., 2012), while in *Phaseolus* annuals and perennials were more closely related (Delgado-Salinas, 2006). Altogether this suggests diverse evolutionary pathways to achieving lifespan differentiation in seed size and germination.

### 4.2 Cultivated accessions show greater seed and plant size relative to wild accessions

Larger seed size in cultivated relative to wild *Lupinus* is consistent with previously documented domestication syndromes in legumes and *Lupinus*. Our results agree with a recent study finding increased seed size in cultivated relative to wild annual Mediterranean species *Lupinus albus* and *L. angustifolius* (Berger et al., 2017). Although *L. albus* was cultivated more than 3000 years before *L. angustifolius*, both species exhibit similar domestication syndromes, including increased seed size, decreased alkaloid content, seed coat softness, and nonshattering fruit, among others (Cowling et al., 1998; Berger et al., 2017). Berger et al. (2017) show that, despite the mean increase in seed size, there is also considerable overlap between cultivated and wild seed size in both *L. albus* and *L. angustifolius*. Our results mirror this for *L. albus*, but we found a greater difference in seed size for cultivated vs. wild *L. angustifolius*, which could be due to differences in sampling. South American *Lupinus mutabilis* is also an ancient domesticate selected for traits such as seed size and nonshattering pods (Atchison et al., 2016). While we could not include the precise perennial wild progenitor (*L. piurensis*) here, its seed size is similar to the other wild perennial *Lupinus* in our study (Atchison et al., 2016). Other perennial domesticated *Lupinus* here (*L. elegans, L. mexicanus, L. rivularis*, and *L. polyphyllus*) contributed less to cultivation differences in seed size, likely due to their different selection history as forages, cover crops, and ornamentals only in the last century (Wolko et al., 2011; Supplementary Table S5).

This study failed to demonstrate any significant changes in germination proportion in cultivated relative to wild *Lupinus* accessions. This was somewhat surprising considering seed coat softness is a trait considered to be part of the domestication syndrome of *L. albus, L. angustifolius*, and *L. mutabilis* (Wolko et al., 2011), likely allowing for loss of physical dormancy. Our scarification treatment may have removed underlying cultivated-wild seed dormancy differences (see discussion above for wild germination). Nevertheless, cultivation did not evidently result in significantly reduced longevity in storage based on germination proportion in *Lupinus*.

*Phaseolus* similarly showed larger seed and vegetative trait values in cultivated species relative to wild species, consistent with domestication syndrome expectations; however, there were some notable differences among annual and perennial comparisons. Consistent with previous studies (Smartt, 1988; Koinange et al., 1996; Aragao et al., 2011), all cultivated annual and perennial *Phaseolus* exhibited greater seed size relative to wild accessions. Perennial *Phaseolus* showed a particularly large difference in cultivated vs. wild seed size, as well as substantial variation in this trait, suggesting a large amount of genetic variation. Much of the perennial seed size variation stemmed from *P. coccineus,* which is primarily outcrossing and more genetically diverse than other *Phaseolus* crops (Bitocchi et al., 2017; Guerra-García et al., 2017). Larger seed size in *P. coccineus* is in agreement with the observations of Delgado-Salinas (1988) and may coincide with a pod-level seed size-number tradeoff, as *P. coccineus* cultivars are also known to have fewer seeds per pod than wild forms. Our seed size observations in USDA accessions are consistent with an analysis of wild and domesticated *Phaseolus* accessions from the International Center for Tropical Agriculture (CIAT), with the exception that they observed larger ranges of seed weight in cultivated annual species (*P. acutifolius, P. vulgaris*; Chacón-Sánchez, 2018). Nevertheless, they similarly found that the perennial *P. coccineus* had the largest range in seed size in both cultivated and wild accessions, that *P. coccineus* showed the largest mean seed weight increase with cultivation, and that the perennial *P. dumosus* also showed some of the largest cultivated seed weights (Chacón-Sánchez, 2018).

Cultivation effects on *Phaseolus* germination may reflect different annual-perennial seed dormancy strategies. Domestication in annual common bean (*P. vulgaris*) and other seed crops has been known to reduce seed coat thickness and seed dormancy (Koinange et al., 1996; Fuller & Allaby, 2009), which could expose cultivated accessions to premature water imbibition and seed mortality in storage (López Herrera et al., 2001). Comparably low germination proportion in cultivated and wild perennial *Phaseolus* could be due to a less selective pressure for dormancy (and therefore lower longevity) in wild perennials relative to wild annuals (see above discussion of wild germination), resulting in less change in germination proportion with domestication. Since our accessions were of various ages and geographic origin, more precise studies on fresh seedstock are necessary to confirm germination differences with cultivation for both annual and perennial species.

Vegetative growth changes with cultivation differed for *Phaseolus* annuals and perennials. Mean node number and stem height were lower in cultivated annual *Phaseolus* and higher in cultivated perennial *Phaseolus* relative to wild accessions. Similarly, node number and height were lower in the cultivated annual *P. vulgaris* (Koinange et al., 1996; Berny Mier y Teran et al., 2018). This suggests that on average cultivated annual *Phaseolus* exhibit lower degrees of stem growth, which may be the result of direct selection for determinate growth or an indirect consequence of increased harvest index, i.e., biomass allocated to reproductive structures at the expense of vegetative growth. This is in spite of all of the cultivated accessions in this study retaining their twining habit. In contrast, cultivated perennial *Phaseolus* exhibited higher values for vegetative growth, possibly indicating a persistence of indeterminate growth in cultivation (Smartt, 1988), allowing for simultaneous selection on increased vegetative growth and reproduction (but see Delgado-Salinas, 1988). Both cultivated annual and cultivated perennial *Phaseolus* displayed higher shoot and root dry mass relative to wild species, in agreement with the findings of Berny Mier y Teran et al. (2018) in *P. vulgaris*; this coincided with higher values in stem diameter for cultivated accessions of both lifespans, in agreement with Smartt (1988). Under cultivation, higher shoot and root biomass relative to wild accessions have also been observed in 30 diverse annual crop species, attributed to greater seed size and leaf area allowing enhanced downstream effects on growth (Milla et al., 2014; Milla & Matesanz, 2017). Our study suggests that some cultivated perennial species in *Phaseolus* exhibit similar size increases.

Root mass fraction, a resource-conservative trait, exhibited higher values in cultivated species relative to wild species for both lifespans; this contradicts the expectation that plants become more resource-acquisitive with domestication (McKey et al., 2012; Vilela & González-Paleo, 2015; Pastor-Pastor et al., 2018). However, higher root mass fraction was also observed in the annuals *Phaseolus vulgaris* (Berny Mier y Teran et al., 2018) and *Pisum sativum* (Weeden, 2007) relative to wild forms. This difference could be a byproduct of a more general higher allocation to vegetative growth, allowing greater amounts of photosynthate to be allocated to roots (Weeden, 2007). This is also consistent with cultivated *P. coccineus* retaining its perennial habit for up to 12 years in tropical climates (Arias, 1980, as cited in Delgado-Salinas, 1988). It is notable that species with the largest seed and vegetative size in this dataset are from tropical environments; there may be additional obstacles in breeding temperate perennial species (or more generally from seasonal environments) given their shorter growing season and their yearly diversion of resources to storage organs to endure an environmental stressor, such as winter. In summary, while many vegetative traits exhibit higher values in cultivated relative to wild accessions for both annual and perennial *Phaseolus* species, some traits may exhibit divergent lifespan patterns. It remains to be determined if this is due to lifespan *per se* or each species’ unique history of artificial selection.

The primary commonality between *Lupinus* and *Phaseolus* was that cultivated species had larger seeds than wild relatives in both annual and perennial comparisons. Larger seed size is a classic domestication syndrome change in annual seed crop domesticates (Purugganan & Fuller, 2009), and our results suggest the same is true for ancient and recent perennial pulse domesticates. Results from *Phaseolus* growth measurements further suggest that cultivated annual and perennial accessions both have greater stem diameter and biomass values relative to wild accessions. Perennial *Phaseolus* species selected primarily for increased seed size and yield may therefore respond in largely the same manner as annual *Phaseolus* domesticates.

### 4.3 Trait covariation is significantly positive in most cases

Positive correlations among traits observed in our dataset suggest that some suites of traits are synergistic, including seed size, some early growth traits, and biomass allocation (**Figure 3**). Positive seed-vegetative size correlations have also been found in other herbaceous plant systems (Geber, 1990; Kleyer et al., 2019), as well as across vascular plants more generally (Díaz et al., 2015). The nearly perfect correlation among seed size parameters (weight, area, and length) suggests that both seed area and mass are jointly selected upon and can be used interchangeably for these species. Significant positive correlations between seed size and vegetative biomass are consistent with the fundamental constraint of plant size on seed size, i.e., small plants cannot produce very large seeds (Venable & Rees, 2009). Furthermore, positive correlations between stem diameter and seed size support the hypothesis that seed size is biomechanically limited by the size of the subtending branch (Aarssen, 2005; Venable & Rees, 2009). While annuals typically have a higher reproductive effort (mass fraction), perennials often are able to produce a greater aboveground biomass in a given season, which may allow for larger seed size and/or total seed yield per plant (Primack, 1979; Dohleman & Long, 2009). Seed size alone may be less reflective of whole-plant reproductive allocation (Vico et al., 2016) and more so of a tolerance-fecundity tradeoff, where larger seeds are produced in fewer numbers but have greater competitive ability in stressful conditions (Muller-Landau, 2010). Nevertheless, the perennial grain crop *Thinopyrum intermedium* showed a positive relationship of total biomass and single seed mass with total reproductive yield across three years of growth, suggesting that these traits are positively correlated in some perennial species (Cattani & Asselin, 2018; also see Kleyer et al., 2019). Lastly, significant positive associations between all dry biomass traits do not support aboveground-belowground tradeoffs; rather, the data suggest that larger plants require greater resource acquisition with proportionally larger roots. This is consistent with previous studies reporting positive correlations among shoot biomass, root biomass, and root mass fraction in *Phaseolus vulgaris* (Berny Mier y Teran et al., 2018).

There were a few notable exceptions to the general trend of positive correlations. There were unexpectedly no significant correlations between seed size and germination proportion. There is a theoretical expectation that germination dormancy and seed bank persistence are negatively related to seed size, since dormancy and seed size entail different strategies of surviving in different environmental conditions, i.e., small, dormant seeds dominate seasonal environments and large, non-dormant seeds are typical of aseasonal environments (Baskin & Baskin, 2014; Rubio de Casas et al., 2017). Instead, the only subset of our data which showed significant germination correlations with seed size, *Lupinus*, exhibited a significant positive relationship between seed size and germination proportion (Supplementary Figure S1E), and the largest-seeded species in this subset inhabit seasonal Mediterranean habitats (except *L. mutabilis*). Our general lack of germination-based correlations may be attributable to our removal of the physical dormancy barrier through scarification, although we may still expect differences in seed longevity in storage proportional to seed size. Other studies have also found that seed size was not significantly correlated with germination rate and proportion (Shipley & Parent, 1991), including in legumes (Nakamura, 1988; Barak et al., 2018).

There was also a lack of significant covariation between early vegetative stem growth traits (stem height and node number) and most biomass traits. This is in contrast to a previous study reporting that stem height is a highly interconnected hub trait in herbaceous perennial species (Kleyer et al., 2019), although we find many of the same positive associations between vegetative and seed size traits. The negative correlation detected in annual species between seed size and node number (Supplementary Figure S1A) may reflect tradeoffs between seed size and stem length allocation. A thicker stem diameter (also negatively correlated with node number; Supplementary Figure S1A) may be required to support the weight of larger seeds, which could come at the expense of stem length. This seed size-node relationship was significantly positive for perennial species (Supplementary Figure S1B), but further research is necessary to determine if such vegetative-reproductive correlations are consistently different between annuals and perennials.

Overall, positive correlations among traits measured here suggests that future breeding efforts targeting greater seed size in perennials may be possible without concomitant reductions in vegetative allocation. Consistently high correlations may also allow for measurement of simpler traits (e.g., stem diameter, shoot dry mass) as proxies for traits that are more difficult to assess (e.g., root dry mass, total yield).

## 4.4 Conclusion

For this set of taxa, we found that cultivation is associated with an increase in seed size and overall vegetative size in both annual and perennial species, and that seed size and most vegetative traits were positively correlated in both cultivated and wild annuals and perennials. While some annual-perennial differences in germination were broadly consistent, lifespan-related seed size differences may be unique to at least the genus level. Thus, lifespan evolution and divergence may take different forms in different lineages, even within the same plant family. The diverse biomes in which each set of annual and perennial species in our study evolved also suggests multiple environmental drivers of life history variation. Traits are shaped and predicted by more specific factors than lifespan alone, including habitat differences, evolutionary history, and which specific organs were targeted during artificial selection. Resolving these factors will require a focused sample of close sister species from the same native range, or intraspecific ecotypes of differing lifespan. Also, all perennials in our study were analyzed in their first year of growth, while their overall life history strategy is contingent upon their survival across multiple years. Thus, this study may also be augmented by multiyear assessments of reproductive and vegetative trait variation in perennial individuals. Such studies will advance our basic knowledge of life history evolution and inform plant breeding as we determine viable methods of ecological intensification. Conclusively, we highlight here common features in cultivated annual and perennial species compared to their wild relatives, and we observed few tradeoffs among seed and vegetative traits in the first year of growth. This offers insight into how perennial legume crops may respond to artificial selection relative to their annual relatives, and it suggests that many traits of interest may be selected for in concert.

## DATA ACCESSIBILITY

The total accession dataset, individual plant dataset, and ImageJ seed analysis dataset for this study can be found in the FigShare online repository under [link to be generated after submission].

## AUTHOR CONTRIBUTIONS

AM and SH designed the project; SH implemented the project and wrote the manuscript, with significant input from AM and MR. MR assisted in statistical analysis and figure generation. CC, TC, and DV provided extensive technical assistance, helpful feedback on the basis and design of the study, and revisions to the manuscript. All authors read and approved the final version of the manuscript.

## ACKNOWLEDGMENTS

The Land Institute provided research tools and space for this project. Seeds were provided by the United States Department of Agriculture Western Regional PI Station (Pullman, WA). We thank the following Land Institute researchers and associates for valuable input in experimental design and intellectual contributions: Lee R. DeHaan, Matthew Newell, Damian A. Ravetta, Alejandra Vilela, and Shuwen Wang. We extend a special thanks to The Land Institute interns, technicians, and associates who assisted in experimental set-up, tending plants, and measuring traits: Mindelena Adams, Sheila Cox, Eliot Cusick, Tiffany Durr, Nick Feijen, Katherine Fortin, Maya Kathrineberg, Laura Kemp, Ron Kinkelar, Jordan Lowry, Maged Nosshi, Diego Sanchez, Codie Van de Meter, and Eline Van de Ven. Brandon Schlautman and REU student Dahlia Martinez also assisted in phenotyping and plant caretaking. We are grateful to the Miller Lab Group for valuable comments on previous versions of this manuscript.

(Figure legends and tables at end of manuscript)

---

**Figure S1.**
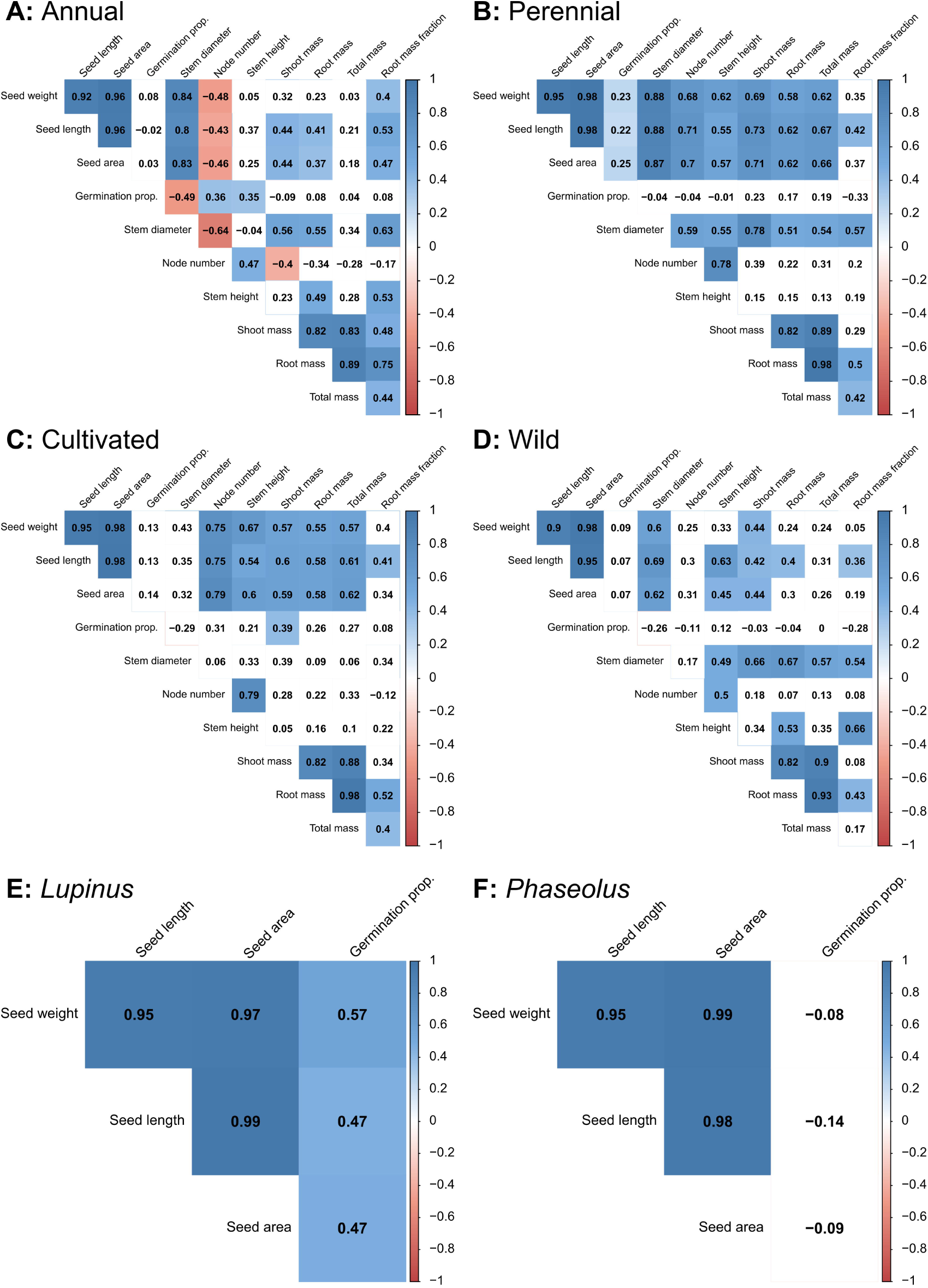
Correlation diagrams of traits for subsets of the dataset: **(A)** annual accessions, **(B)** perennial accessions, **(C)** wild accessions, **(D)** cultivated accessions, **(E)** *Lupinus* accessions (seed traits only), and **(F)** *Phaseolus* accessions (seed traits only). Each category includes all other data which match the criterion (e.g., ‘annual’ includes both cultivated and wild, *Lupinus* and *Phaseolus* annuals). Numbers in boxes represent the Pearson correlation coefficient. Blue and red colors indicate significant positive and negative correlations (at the α = 0.05 level), respectively; absence of color signifies lack of significance.

