## Supplemental Table S1 for "Comparative analysis of seed size, germination, and vegetative allocation in annual and herbaceous perennial crops and their wild relatives in *Lupinus* and *Phaseolus* (Fabaceae)"

**Table S1.** Mean seed and vegetative trait values, with one standard deviation, grouped by genus, lifespan, and cultivation status. Different letters indicate a significant difference (at least *P* < 0.05) within a trait between lifespan × cultivation groups according to a *post-hoc* Tukey HSD test run on the linear model, with covariates included.

| Genus |  | Group | | | |
| --- | --- | --- | --- | --- | --- |
| *Lupinus* | Trait | cultivated annual | wild annual | cultivated perennial | wild perennial |
|  | Single seed weight (mg) | 256.91 ± 95.54 ^B^ | 140.20 ± 113.36 ^A^ | 88.16 ± 2.45 ^D^ | 21.04 ± 8.32 ^C^ |
|  | Single seed length (mm) | 9.64 ± 1.62 ^B^ | 7.16 ± 2.63 ^A^ | 6.96 ± 2.73 ^A^ | 4.50 ± 0.75 ^C^ |
|  | Single seed area (mm^2^) | 58.76 ± 20.79 ^B^ | 34.50 ± 25.75 ^A^ | 32.80 ± 24.56 ^A^ | 10.18 ± 3.84 ^C^ |
|  | Germination proportion | 0.87 ± 0.25 ^A^ | 0.88 ± 0.13 ^A^ | 0.44 ± 0.23 ^B^ | 0.37 ± 0.28 ^B^ |
| *Phaseolus* | Trait | cultivated annual | wild annual | cultivated perennial | wild perennial |
|  | Single seed weight (mg) | 152.83 ± 67.96 ^A^ | 59.24 ± 55.94 ^A^ | 646.15 ± 313.99 ^B^ | 101.11 ± 97.78 ^A^ |
|  | Single seed length (mm) | 8.58 ± 1.65 ^B^ | 6.14 ± 2.08 ^A^ | 15.12 ± 3.15 ^C^ | 7.33 ± 2.09 ^AB^ |
|  | Single seed area (mm^2^) | 38.45 ± 11.82 ^A^ | 20.19 ± 12.58 ^A^ | 128.66 ± 49.04 ^B^ | 31.27 ± 19.67 ^A^ |
|  | Germination proportion | 0.55 ± 0.34 ^B^ | 0.92 ± 0.14 ^A^ | 0.57 ± 0.34 ^B^ | 0.55 ± 0.41 ^B^ |
|  | Stem diameter (mm) | 2.30 ± 0.74 ^B^ | 1.24 ± 0.35 ^A^ | 2.53 ± 0.79 ^B^ | 1.16 ± 0.45 ^A^ |
|  | Node number | 2.66 ± 0.77 ^A^ | 3.53 ± 0.50 ^A^ | 4.46 ± 1.07 ^B^ | 3.46 ± 0.78 ^AB^ |
|  | Stem height (cm) | 23.76 ± 13.66 ^A^ | 31.94 ± 16.83 ^A^ | 30.37 ± 19.78 ^A^ | 22.38 ± 14.74 ^A^ |
|  | Shoot dry mass (g) | 1.06 ± 0.46 ^BC^ | 0.73 ± 0.36 ^A^ | 2.04 ± 1.06 ^C^ | 0.55 ± 0.57 ^AB^ |
|  | Root dry mass (g) | 0.47 ± 0.40 ^A^ | 0.32 ± 0.24 ^A^ | 1.60 ± 1.36 ^B^ | 0.24 ± 0.28 ^A^ |
|  | Total dry mass (g) | 1.67 ± 0.89 ^A^ | 1.48 ± 0.84 ^A^ | 5.19 ± 3.66 ^B^ | 0.99 ± 1.03 ^A^ |
|  | Root mass fraction | 0.25 ± 0.10 ^AB^ | 0.22 ± 0.07 ^A^ | 0.29 ± 0.07 ^B^ | 0.22 ± 0.10 ^AB^ |
