## Supplemental Table S2 for "Comparative analysis of seed size, germination, and vegetative allocation in annual and herbaceous perennial crops and their wild relatives in *Lupinus* and *Phaseolus* (Fabaceae)"

**Table S2.** Results of linear models used to assess broad geographic effects on lifespan-related seed trait variation in wild *Lupinus* and wild *Phaseolus*. Letters denote separate models with different covariates run independently for each genus: (a, c) seed size traits and (b, d) germination.

| *Lupinus* | Trait | Geography | Lifespan† | Geography *×* Lifespan† | Species |  |  |  |  |
| --- | --- | --- | --- | --- | --- | --- | --- | --- | --- |
| (a) | Seed weight | *F*_1_ = 22.39*** | *F*_1_ = 0.28 | ---- | *F*_6_ = 24.88*** |  |  |  |  |
|  | Seed area | *F*_1_ = 42.11*** | *F*_1_ = 1.65 | ---- | *F*_6_ = 49.92*** |  |  |  |  |
|  | Seed length | *F*_1_ = 46.52*** | *F*_1_ = 6.72* | ---- | *F*_6_ = 23.59*** | Age | Scarification | Soak time | Seed damage |
| (b) | Germination probability | *F*_1_ = 0.04 | *F*_1_ = 4.44 | ---- | *F*_6_ = 2.22 | *F*_1_ = 1.25 | *F*_1_ = 1.10 | *F*_1_ = 0.74 | *F*_1_ = 0.06 |
| *Phaseolus* | Trait | Geography | Lifespan | Geography *×* Lifespan | Species |  |  |  |  |
| (c) | Seed weight | *F*_1_ = 13.16*** | *F*_1_ = 1.71 | *F*_1_ = 0.24 | *F*_3_ = 0.92 |  |  |  |  |
|  | Seed area | *F*_1_ = 24.00*** | *F*_1_ = 2.95 | *F*_1_ = 0.91 | *F*_3_ = 0.52 |  |  |  |  |
|  | Seed length | *F*_1_ = 36.22*** | *F*_1_ = 1.39 | *F*_1_ = 0.68 | *F*_3_ = 1.48 | Age | Scarification†† | Soak time | Seed damage |
| (d) | Germination probability | *F*_1_ = 6.18* | *F*_1_ = 11.91** | *F*_1_ = 11.86** | *F*_3_ = 0.50 | *F*_1_ = 1.44 | ---- | *F*_1_ = 0.37 | *F*_1_ = 1.47 |

**P* < 0.05; ***P* < 0.01; ****P* < 0.001.

† Geography *×* Lifespan cannot be assessed for *Lupinus* due to the absence of perennial Mediterranean species.

†† Scarification treatment was the same for all *Phaseolus* accessions.
