## Supplemental Table S3 for "Comparative analysis of seed size, germination, and vegetative allocation in annual and herbaceous perennial crops and their wild relatives in *Lupinus* and *Phaseolus* (Fabaceae)"

**Table S3.** Results of linear models used to assess broad geographic effects on lifespan-related vegetative trait variation in wild *Phaseolus*. Letters denote separate models with different covariates, while the main effects are the same for all traits: (a) early vegetative growth traits and (b) biomass traits.

|  | Trait | Geography | Lifespan | Geography *×* Lifespan | Species | Health |  |  |
| --- | --- | --- | --- | --- | --- | --- | --- | --- |
| (a) | Stem diameter | *F*_1_ = 17.10*** | *F*_1_ = 0.72 | *F*_1_ = 1.19 | *F*_2_ = 0.72 | *F*_1_ = 0.50 |  |  |
|  | Node number | *F*_1_ = 0.02 | *F*_1_ = 0.10 | *F*_1_ = 7.75** | *F*_2_ = 1.46 | *F*_1_ = 6.51* |  |  |
|  | Stem height | *F*_1_ = 8.60** | *F*_1_ = 1.95 | *F*_1_ = 0.85 | *F*_2_ = 3.58* | *F*_1_ = 1.83 |  |  |
|  | Trait | Geography | Lifespan | Geography *×* Lifespan | Species | Health | Reproductive state | Outdoor proportion |
| (b) | Shoot dry mass | *F*_1_ = 6.16* | *F*_1_ = 1.74 | *F*_1_ = 1.84 | *F*_2_ = 0.85 | *F*_1_ = 2.03 | *F*_1_ = 0.35 | *F*_1_ = 0.00 |
|  | Root dry mass | *F*_1_ = 10.00** | *F*_1_ = 1.51 | *F*_1_ = 0.04 | *F*_2_ = 0.31 | *F*_1_ = 1.05 | *F*_1_ = 0.04 | *F*_1_ = 2.03 |
|  | Total dry mass | *F*_1_ = 4.98* | *F*_1_ = 2.42 | *F*_1_ = 1.11 | *F*_2_ = 0.18 | *F*_1_ = 1.26 | *F*_1_ = 0.01 | *F*_1_ = 0.55 |
|  | Root mass fraction | *F*_1_ = 15.94*** | *F*_1_ = 0.01 | *F*_1_ = 2.67 | *F*_2_ = 0.73 | *F*_1_ = 2.02 | *F*_1_ = 0.50 | *F*_1_ = 2.91 |

* *P* < 0.05; ** *P* < 0.01; *** *P* < 0.001.
