## Supplemental Table S4 for "Comparative analysis of seed size, germination, and vegetative allocation in annual and herbaceous perennial crops and their wild relatives in *Lupinus* and *Phaseolus* (Fabaceae)"

**Table S4.** Mean seed and vegetative trait values, with one standard deviation, grouped by lifespan and broad geographic distribution for wild *Lupinus* and *Phaseolus*. Different letters indicate a significant difference (at least *P* < 0.05) within a trait between lifespan × geography groups according to a *post-hoc* Tukey HSD test run on the linear model, with covariates included.

| Genus |  | Group | | | |
| --- | --- | --- | --- | --- | --- |
| *Lupinus* | Trait | Mediterranean annual† | American perennial† |  |  |
|  | Single seed weight (mg) | 150.61 ± 110.80 ^A^ | 21.04 ± 8.32 ^B^ |  |  |
|  | Single seed length (mm) | 7.51 ± 2.38 ^A^ | 4.50 ± 0.75 ^B^ |  |  |
|  | Single seed area (mm^2^) | 36.87 ± 25.15 ^A^ | 10.18 ± 3.84 ^B^ |  |  |
|  | Germination proportion | 0.88 ± 0.14 ^A^ | 0.37 ± 0.28 ^B^†† |  |  |
| *Phaseolus* | Trait | desert annual | desert perennial | tropical annual | tropical perennial |
|  | Single seed weight (mg) | 20.89 ± 14.83 ^A^ | 32.94 ± 18.36 ^AB^ | 89.92 ± 58.05 ^AB^ | 123.83 ± 103.28 ^B^ |
|  | Single seed length (mm) | 4.47 ± 1.10 ^A^ | 5.19 ± 1.33 ^A^ | 7.59 ± 1.58 ^B^ | 8.04 ± 1.81 ^B^ |
|  | Single seed area (mm^2^) | 11.06 ± 5.74 ^A^ | 14.56 ± 7.32 ^AB^ | 28.11 ± 11.51 ^B^ | 36.84 ± 19.41 ^B^ |
|  | Germination proportion | 0.86 ± 0.17 ^A^ | 1.00 ± 0.00 ^B^ | 0.98 ± 0.04 ^C^ | 0.40 ± 0.36 ^D^ |
|  | Stem diameter (mm) | 1.05 ± 0.15 ^A^ | 0.84 ± 0.23 ^A^ | 1.44 ± 0.38 ^B^ | 1.49 ± 0.36 ^B^ |
|  | Node number | 3.56 ± 0.44 ^A^ | 3.00 ± 0.86 ^A^ | 3.49 ± 0.58 ^A^ | 3.93 ± 0.32 ^A^ |
|  | Stem height (cm) | 23.49 ± 12.99 ^A^ | 9.37 ± 3.29 ^A^ | 45.06 ± 13.63 ^B^ | 35.38 ± 6.73 ^AB^ |
|  | Shoot dry mass (g) | 0.66 ± 0.33 ^A^ | 0.18 ± 0.14 ^A^ | 0.81 ± 0.39 ^A^ | 0.86 ± 0.61 ^A^ |
|  | Root dry mass (g) | 0.21 ± 0.14 ^A^ | 0.06 ± 0.06 ^A^ | 0.45 ± 0.26 ^A^ | 0.36 ± 0.31 ^A^ |
|  | Total dry mass (g) | 1.26 ± 0.59 ^A^ | 0.26 ± 0.17 ^A^ | 1.68 ± 1.00 ^A^ | 1.47 ± 1.09 ^A^ |
|  | Root mass fraction | 0.16 ± 0.05 ^A^ | 0.20 ± 0.09 ^AB^ | 0.27 ± 0.05 ^B^ | 0.24 ± 0.10 ^AB^ |

† There are no known Mediterranean perennials, and there is only one accession of North American annual in this study (*L. arizonicus*), which could not be used in the Tukey test.

†† Nonsignificant covariates were dropped from the *Lupinus* germination linear model assessed here.
