## Supplemental Table S5 for "Comparative analysis of seed size, germination, and vegetative allocation in annual and herbaceous perennial crops and their wild relatives in *Lupinus* and *Phaseolus* (Fabaceae)"

**Table S5.** Species-level summary of native distribution in their wild form, cultivated uses, and timeline of its earliest estimated cultivation. This represents the overall description of each species and does not necessarily reflect all accessions used in this study. Livestock feed may include either forage and/or seed consumption. Pulse refers to dry legume seeds consumed by humans. Vegetable refers either vegetative parts or green pods consumed by humans. A blank signifies no known cultivated uses (not counting ethnobotanical uses here). For species in which only one cultivated accession was used in this study, the name of the cultivar used is included in quotes.

| **Genus** | **Lifespan** | **Species** | **Native distribution*** | **Cultivated uses** | **Original cultivation** | **Citations** |
| --- | --- | --- | --- | --- | --- | --- |
| *Lupinus* | annual | *L. albus* | Greece to W Turkey (M) | cover crop; livestock feed; pulse | 2000 BCE | Gladstones 1974; Cowling et al. 1998; Kurlovich 2002 |
|  |  | *L. angustifolius* | Pan-Mediterranean (M) | cover crop; livestock feed; pulse | 1860s CE | Cowling et al. 1998; Clements et al. 2005; Wolko et al. 2011 |
|  |  | *L. arizonicus* | SW United States (A) |  |  | USDA-NRCS 2019 |
|  | perennial | *L. albicaulis* | W United States (A) |  |  | USDA-NRCS 2019 |
|  |  | *L. albifrons* | W United States (A) |  |  | USDA-NRCS 2019 |
|  |  | *L. andersonii* | W United States (A) |  |  | USDA-NRCS 2019 |
|  |  | *L. elegans* | Mexico (A) | cover crop (“Armex”) | 1957 CE | Surrency & Undayag 2000; Wolko et al. 2011 |
|  |  | *L. mexicanus* | Mexico (A) | ornamental | 1950s CE | Burkart 1959; Clements et al. 2005; Wolko et al. 2011 |
|  |  | *L. mutabilis* | N Peru (A)** | pulse; vegetable | 600 BCE | Gross 1986; Cowling et al. 1998; Wolko et al. 2011; Atchison et al. 2016 |
|  |  | *L. polyphyllus* | W North America (A) | cover crop; livestock feed; ornamental | 1930s CE | Wolko et al. 2011; Beuthin 2012; Kurlovich & Heinänen 2002; Clements et al. 2005; USDA-NRCS 2019 |
|  |  | *L. rivularis* | NW United States (A) | cover crop; ornamental (“Hederma”) | 1969 CE | USDA-NRCS 2012, 2019 |
| **Genus** | **Lifespan** | **Species** | **Native distribution** | **Cultivated uses** | **Original cultivation**† | **Citations** |
| *Phaseolus* | annual | *P. acutifolius* | SW United States to Mexico (D) | pulse | 500 BCE | Buhrow 1983; Smartt 1988; Kaplan & Lynch 1999; Bitocchi et al. 2017 |
|  |  | *P. filiformis* | SW United States to Mexico (D) |  |  | Buhrow 1983 |
|  |  | *P. vulgaris* | W Mesoamerica and W South America (T) | pulse; vegetable | 3030 BCE | Smartt 1988; Kaplan & Lynch 1999; Bitocchi et al. 2017 |
|  | perennial | *P. angustissimus* | SW United States to Mexico (D) |  |  | Buhrow 1983 |
|  |  | *P. coccineus* | Mexico to Panama (T) | pulse; vegetable | 880 CE | Smartt 1976; Delgado-Salinas 1988; Smartt 1988; Kaplan & Lynch 1999 |
|  |  | *P. dumosus* | Guatemala (T) | pulse; vegetable | †† | Smartt 1988; Schmit & Debouck 1991; |
|  |  | *P. maculatus* | SW United States to Mexico (D) |  |  | Buhrow 1983 |

* Capital letters denote the broad geographic distribution assigned to each species that was used in linear models: For *Lupinus*, M refers to Mediterranean annuals and A to American perennials (except the annual *L. arizonicus*); For *Phaseolus*, D refers to desert and T to tropical.

** Based on *Lupinus piurensis*, the most likely wild progenitor of *L. mutabilis* (Atchison et al. 2016).

† Evidence for the earliest cultivation in *Phaseolus* species is generally highly variable, ranging from 9000 BCE to less than 500 years ago; here we use more recent radiocarbon dates taken directly from *Phaseolus* material, but we recognize that this is dependent upon material that was carbonized or did not decay significantly (Kaplan & Lynch 1999).

†† To our knowledge, the timing and precise nature of *Phaseolus dumosus* domestication has not yet been determined.
